# *Lotus japonicus* symbiosis genes impact microbial interactions between symbionts and multikingdom commensal communities

**DOI:** 10.1101/547687

**Authors:** Thorsten Thiergart, Rafal Zgadzaj, Zoltán Bozsóki, Ruben Garrido-Oter, Simona Radutoiu, Paul Schulze-Lefert

**Affiliations:** Max Planck Institute for Plant Breeding Research, 50829 Cologne, Germany; Cluster of Excellence on Plant Sciences (CEPLAS), Max Planck Institute for Plant Breeding Research, 50829 Cologne, Germany; Department of Molecular Biology and Genetics, Faculty of Science and Technology, Aarhus University, 8000 C Aarhus, Denmark

## Abstract

The legume *Lotus japonicus* engages in mutualistic symbiotic relationships with arbuscular mycorrhiza (AM) fungi and nitrogen-fixing rhizobia. Using plants grown in natural soil and community profiling of bacterial 16S *rRNA* genes and fungal internal transcribed spacers (ITS), we examined the role of the *Lotus* symbiosis genes *RAM1, NFR5, SYMRK*, and *CCaMK* in structuring bacterial and fungal root-associated communities. We found host genotype-dependent community shifts in the root and rhizosphere compartments that were confined to bacteria in *nfr5* or fungi in *ram1* mutants, whilst *symrk* and *ccamk* plants displayed changes across both microbial kingdoms. We observed in all AM mutant roots an almost complete depletion of Glomeromycota taxa that was accompanied by a concomitant enrichment of Helotiales and Nectriaceae fungi, suggesting compensatory niche replacement within the fungal community. A subset of Glomeromycota whose colonization is strictly dependent on the common symbiosis pathway was retained in *ram1* mutants, indicating that *RAM1* is dispensable for intraradical colonization by some Glomeromycoyta fungi. However, intraradical colonization by bacteria belonging to the Burkholderiaceae and Anaeroplasmataceae is dependent on AM root infection, revealing a microbial interkingdom interaction. Despite an overall robustness of the bacterial root microbiota against changes in the composition of root-associated fungal assemblages, bacterial and fungal co-occurrence network analysis demonstrates that simultaneous disruption of AM and rhizobia symbiosis increases the connectivity among taxa of the bacterial root microbiota. Our findings imply a broad role for *Lotus* symbiosis genes in structuring the root microbiota and identify unexpected microbial interkingdom interactions between root symbionts and commensal communities.

**Importance:** Studies on symbiosis genes in plants typically focus on binary interactions between roots and soil-borne nitrogen-fixing rhizobia or mycorrhizal fungi in laboratory environments. We utilized wild-type and symbiosis mutants of a model legume, grown in natural soil, in which the bacterial or fungal or both symbioses are impaired to examine potential interactions between the symbionts and commensal microorganisms of the root microbiota when grown in natural soil. This revealed microbial interkingdom interactions between the root symbionts and fungal as well as bacterial commensal communities. Nevertheless, the bacterial root microbiota remains largely robust when the fungal symbiosis is impaired. Our work implies a broad role for host symbiosis genes in structuring the root microbiota of legumes.

## Introduction

Mutualistic plant-microbe interactions are essential adaptive responses dating back to plant colonization of terrestrial habitats [1,2]. Endosymbiotic association with obligate arbuscular mycorrhizal (AM) fungi belonging to the phylum Glomeromycota is considered to have enabled early land plants to adapt to and survive harsh edaphic conditions by improving the acquisition of nutrients, especially phosphorus, from soil [3]. It is estimated that approximately 80% of extant plant species remain proficient in AM symbiosis (AMS), testifying to its importance for survival in natural ecosystems [4,5,6]. Another more recent endosymbiotic relationship has evolved between plants belonging to distinct lineages of flowering plants (Fabales, Fagales, Cucurbitales, and Rosales) and nitrogen-fixing members of the Burkholderiales, Rhizobiales or Actinomycetales, enabling survival on nitrogen-poor soils. These bacteria fix atmospheric nitrogen under the low oxygen conditions that are provided by plant root nodules.

Studies using mutant legumes deficient in both AM and root nodule symbiosis (RNS) revealed that a set of genes defined as the common symbiotic signaling pathway (CSSP) are crucial for these symbioses. In the model legume *Lotus japonicus*, Nod factor perception by NFR1 and NFR5 activates downstream signaling through SYMRK, a malectin and leucine-rich repeat (LRR)-containing receptor-like kinase (RLK) [7], currently considered to be the first component of the CSSP. SYMRK associates with NFR5 through a mechanism involving intramolecular cleavage of the SYMRK ectodomain, thereby exposing its LRR domains [8]. Signaling from the plasma membrane is transduced to the nuclear envelope where ion channels [9,10], nuclear pore proteins [11,12,13] and cyclic nucleotide-gated channels [14] mediate symbiotic calcium oscillations. These calcium oscillations are interpreted by the calcium- and calmodulin-dependent protein kinase CCaMK, [15,16] that interacts with the DNA binding transcriptional activator CYCLOPS [17,18,19]. Several GRAS transcription factors (NSP1, NSP2, RAM1, RAD1) are activated downstream of CCaMK and CYCLOPS and determine whether plants engage in AMS or RNS symbiosis.

Plants establish symbioses with AM fungi and nitrogen-fixing bacteria by selecting interacting partners from the taxonomically diverse soil biome. These interactions are driven by low mineral nutrient availability in soil and induce major changes in host and microbial symbiont metabolism [20,21]. Although RNS develops as localized events on legume roots, analysis of *Lotus* mutants impaired in their ability to engage in symbiosis with nitrogen-fixing bacteria revealed that these mutations do not only abrogate RNS, but also impact the composition of taxonomically diverse root- and rhizosphere-associated bacterial communities, indicating an effect on multiple bacterial taxa that actively associate with the legume host, irrespective of their symbiotic capacity [22]. By contrast, the effect of AMS is known to extend outside the host *via* a hyphal network that can penetrate the surrounding soil and even indirectly affect adjacent plants [23]. In soil, fungal hyphae themselves represent environmental niches and are populated by a specific set of microbes [24]. Although the biology of AMF is well understood and genetic disruption of AMS was recently shown to exert a relatively small effect on root-associated fungal communities in *Lotus* [25], the potential impact of AMS and/or RNS on root-associated bacterial and fungal commensals remains poorly understood, mainly because previous studies have their focus on either bacteria [22] or fungi alone [25].

We reasoned that the model legume *Lotus japonicus*, with its well-characterized symbiosis signaling mutants impaired in RNS, AMS, or both is particularly useful to examine whether genetic perturbations of these symbioses impact only commensal communities of the corresponding microbial kingdom and/or influence microbial interkingdom interactions in the root microbiota. We applied bacterial and fungal community profiling experiments to root samples collected from wild-type (WT) *L. japonicus* and four symbiosis signaling mutants, grown in natural soil. We show that genetic disruption of the symbioses results in significant host genotype-dependent microbial community shifts in the root and surrounding rhizosphere compartments. These changes were mainly confined to either bacterial or fungal communities in RNS- or AMS-deficient plant lines, respectively, whereas mutants with defects in the CSPP revealed major changes in assemblages of the root microbiota across both microbial kingdoms. We found that perturbation of AM symbiosis alone is sufficient to deplete a subset of bacterial taxa belonging to the Burkholderiaceae and Anaeroplasmataceae families from the root microbial community, whereas simultaneous perturbation of AM and rhizobia symbioses increases the connectivity within the bacterial root co-occurrence network.

## Results

### Root fractionation protocol affects the composition of associated bacterial communities

Earlier physiological studies have shown that only cells of a specific developmental stage, located in the root elongation zone, respond to Myc and Nod factors, mount symbiotic calcium oscillations and enable epidermal infection by rhizosphere-derived fungal and bacterial symbionts [26,27]. To explore spatial organization of root-associated bacterial and fungal communities along the longitudinal axis, we collected samples of the upper and lower root zones as well as the entire root system of 10 week-old Gifu wild-type plants, grown in Cologne soil (2 to 5 cm and >9 cm of the root system, respectively; Fig. 1A; [28]). Microbial assemblages of these three root endosphere compartments were compared to the communities in the corresponding rhizosphere fractions, i.e. soil tightly adhering to the respective root zones, and with the bacterial biome present in unplanted Cologne soil. 16S rRNA gene amplicon libraries of the V5-V7 hypervariable region and gene libraries of the Internally Transcribed Spacer 2 (ITS2) region of the eukaryotic ribosome were generated by amplification [29,30,31]. Information on the number and relative abundance of operational taxonomic units (OTUs) in each compartment was used to calculate α− (Shannon index; within sample diversity) and β-diversity (Bray-Curtis distances; between samples diversity), OTU enrichment and taxonomic composition. In bacteria we observed a gradual decrease in α-diversity from unplanted soil to the rhizosphere and to the root endosphere compartments, a trend that was similar for each longitudinal root fraction. This suggests that winnowing of root commensals from the highly complex soil biome occurs in all tested root zones (Fig. S1A). Similar overall results were obtained for the fungal dataset (Fig. S1B), but the decrease in diversity from unplanted soil towards the rhizosphere was mild or even lacking. The latter finding is similar to that of a recent study of root-associated fungi in non-mycorrhizal *A. thaliana* sampled at three natural sites [32]. Analyses of taxonomic composition and β-diversity revealed striking differences in the endosphere and rhizosphere compartments associated with the upper and lower root longitudinal fractions. The composition of bacterial and fungal taxa of the whole root closely resembled that of the upper root fraction (Fig. 1B), with only low numbers of OTUs differentially abundant between these two compartments (Fig. 1C and 1D). This suggests that microbes colonizing the lower root fraction constitute only a small fraction of the entire *Lotus* root microbiota. Additionally, we observed a higher sample-to-sample variation in the taxonomic profiles of the lower root zone compared to the upper whole root fractions (Fig. 1B). This greater community variation in the developmentally younger region of *L. japonicus* roots might reflect a nascent root microbiota or greater variation in root tissue and adherent rhizosphere samples that we recovered from this root zone by our fractionation protocol. Based on the finding that whole root and upper root compartments host comparable bacterial communities and given their greater stability we decided to use the former for further analyses.

**Fig1.**
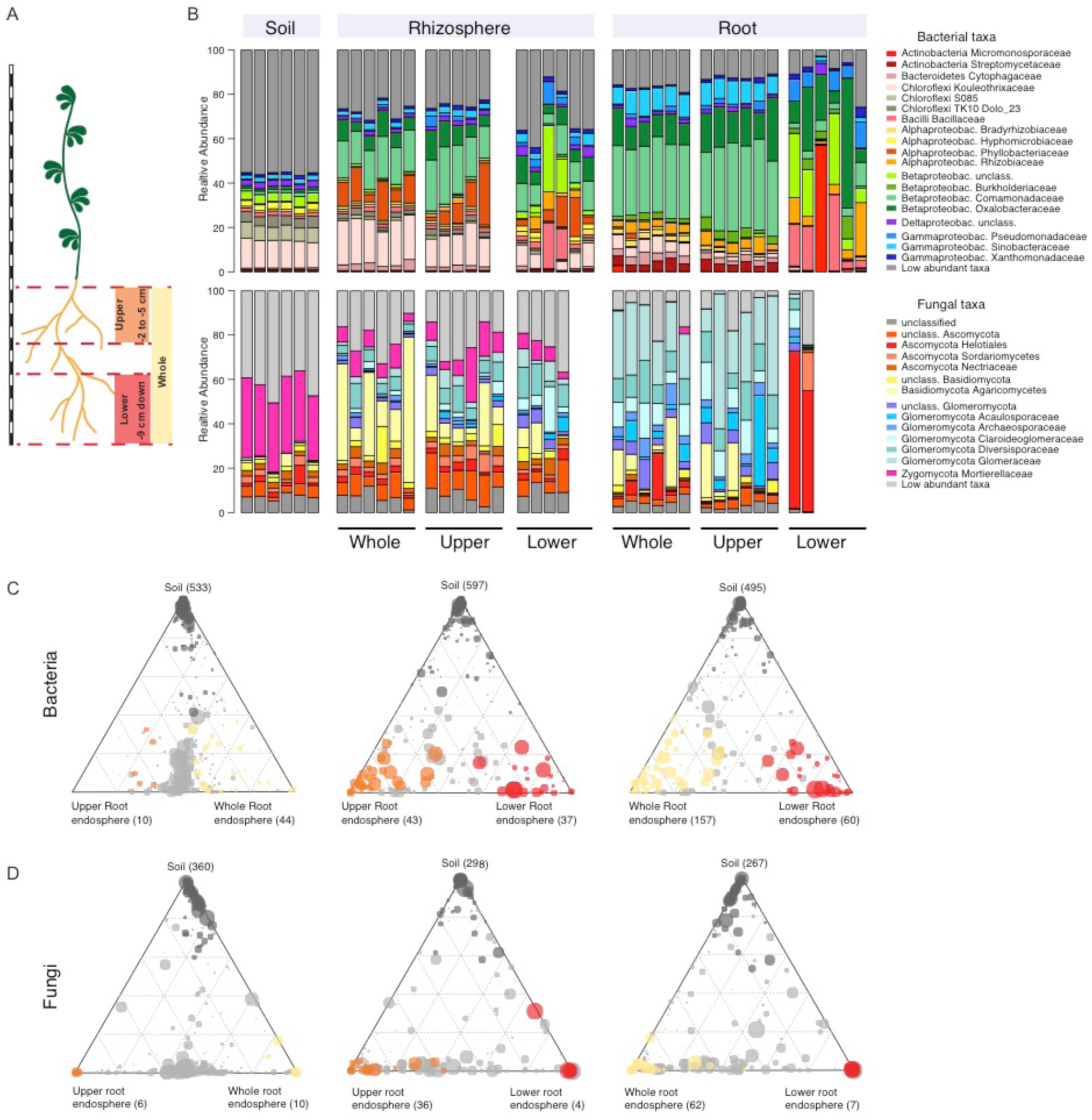
Bacterial & fungal community profile for different root fractions of *L. japonicus*. A) Cartoon showing the length of the three different root fractions. B) Community profile showing the relative abundance of bacterial (upper panel) and fungal (lower panel) families across compartments and fractions (only samples with >5000 (bacteria) or >1000 (fungi) reads are shown, taxa having average RA < 0.1 (bacteria) or <0.15 (fungi) across all samples are aggregated as low-abundant.). C) Ternary plots showing bacterial OTUs that are enriched in the endosphere of specific root fractions, compared to the soil samples. B) Ternary plots showing fungal OTUs that are enriched in the endosphere of specific root fractions, compared to the soil samples. Circle size corresponds to RA across all fractions. Dark grey circles denote OTUs that are enriched in soil, light grey circles always represent OTUs that are not enriched in any of the fractions.

### Host genes needed for symbioses determine bacterial and fungal community composition of *L. japonicus* root and rhizosphere

For root microbiota analysis, we cultivated wild-type (ecotype Gifu) *L. japonicus* and *nfr5-2, symrk-3, ccamk-13* and *ram1-2 (nfr5, symrk, ccamk* and *ram1*, from thereof) mutant genotypes in parallel in two batches of Cologne soil, to account for batch-to-batch and seasonal variation at the sampling site. *nfr5-2* mutant plants are impaired in rhizobial Nod factor perception and signaling, which prevents initiation of infection thread formation [33]. Mutations in *SymRK* and *CCaMK* affect the common symbiosis pathway downstream of Nod or Myc factor perception, abrogating infection either by nitrogen fixing rhizobia or AM fungi [7,34]. The RAM1 transcription factor controls arbuscule formation, and while *ram1* mutants of *L. japonicus* are indistinguishable from wild type and permit incipient AM fungus infection, fungal colonization is terminated with the formation of stunted symbiotic structures [35]. All plant genotypes appeared healthy (Fig. 2A-E), but the shoot length and shoot fresh weight of all mutant plants was significantly reduced in comparison to wild type (Fig. 2F and 2G), suggesting that genetic disruption of either AM or *Rhizobium* symbiosis is detrimental for the fitness of plants grown in natural soil. All genetic defects in nitrogen-fixing symbiosis, validated by the absence of root nodules in *nfr5, symrk* and *ccamk* genotypes (Fig. 2C-E, Table S1), resulted in similarly severe impacts on plant growth (Fig. 2F and 2G), whereas both shoot length and shoot fresh weight were significantly but less severely reduced in *ram1* plants. *ram1* plants still formed nodules and, unlike WT and *nfr5*, showed impairment in AM symbiosis (Table S1).

**Fig2.**
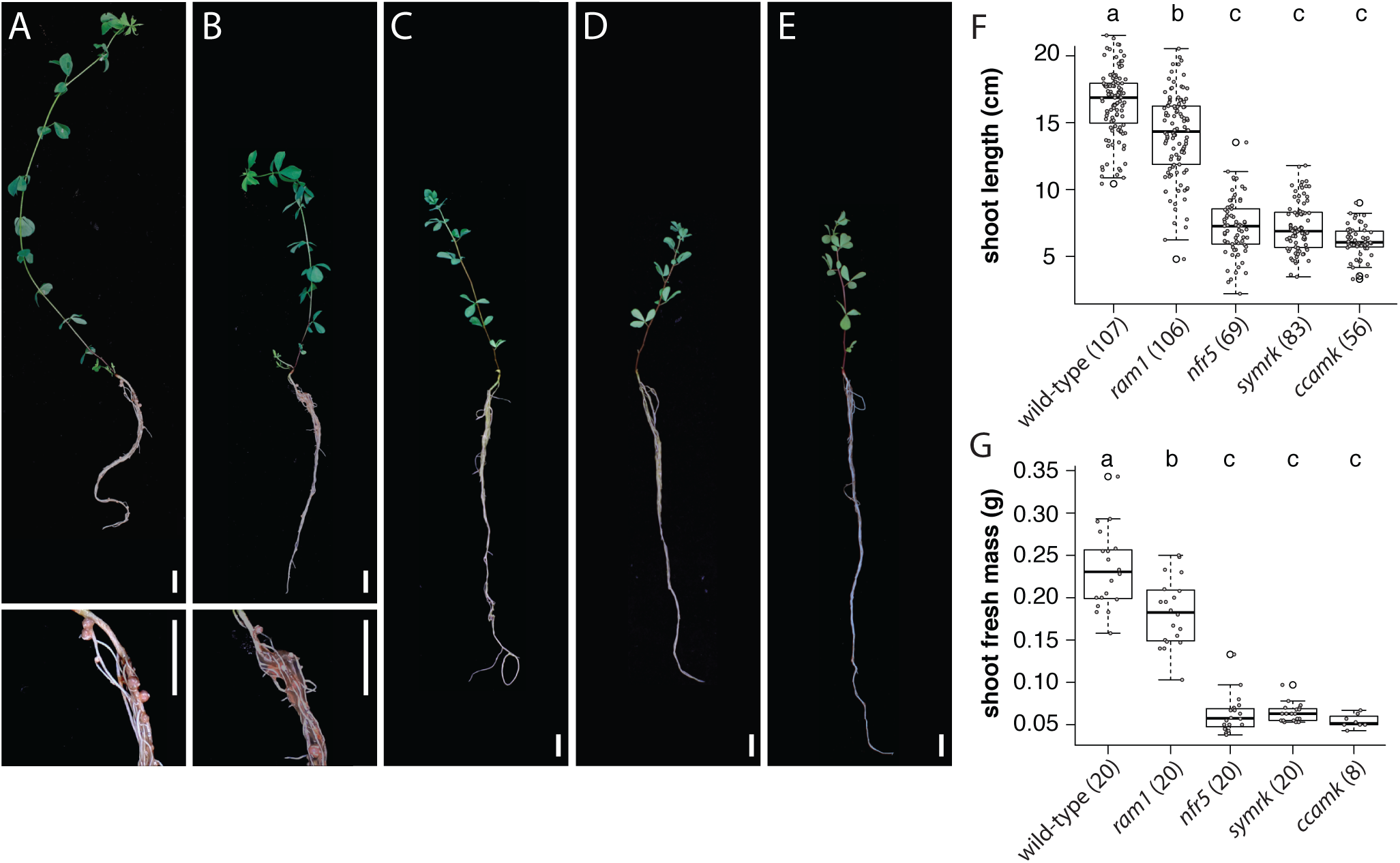
Phenotypes of WT and mutant plants. Images depicting *L. japonicus* wild-type (A) and symbiosis-deficient mutant plants: *ram1* (B), *nfr5* (C), *symrk* (D) and *ccamk* (E). Insets show close-up view of nodules. Scale bars correspond to 1 cm. F) Boxplots display the shoot length for the identical set of genotypes presented in (A-E). G) Boxplots displaying the shoot fresh mass. Letters above plots correspond to groups based on Tukey’s HSD test (P<0.05). Number of samples are indicated in brackets.

In order to determine the impact of rhizobial and AM symbiosis on root microbiota assembly, we characterized fungal and bacterial communities of unplanted Cologne soil, rhizosphere, and root compartments of all aforementioned *L. japonicus* genotypes at bolting stage. Visible nodules and root primordia were removed from the roots of nodulating wild type and *ram1* genotypes prior to sample processing for community profiling. We amplified the V5-V7 hypervariable region of the bacterial 16S rRNA gene and the ITS2 region of the eukaryotic ribosomal genes. High-throughput sequencing of these amplicons yielded 22,761,657 16S and 21,228,781 ITS reads, distributed in 222 and 274 samples, respectively, which were classified into 5,780 and 3,361 distinct microbial OTUs. Analysis of α-diversity revealed a general reduction of complexity from unplanted soil to rhizosphere and lastly to root compartments for bacterial communities, whereas the complexity of fungal communities was similar for the plant-associated compartments (Fig. S2A and S2B), which is consistent with a recent study of *A. thaliana* root-associated fungal communities [32]. Bacterial α-diversity was slightly elevated in the *nfr5* genotype in rhizosphere and root compartments in comparison to all other genotypes (Fig. S2A). Fungal communities were similarly diverse in the rhizosphere of all tested plant genotypes, but their diversity in the root compartment was significantly and specifically reduced in all three AM mutants (*ccamk, ram1*, and *symrk*; Fig. S2B).

Analysis of β-diversity using Principal Coordinate Analysis (PCoA) of Bray-Curtis distances showed a significant effect of soil batch on soil-resident bacterial and fungal communities (Fig. S2C and D). In order to account for this technical factor and assess the impact of the different host compartment and genotypes in community composition, we performed a Canonical Analysis of Principle Components Coordinates (CAP; [36]). This revealed a clear differentiation of bacterial and fungal communities in the tested plant genotypes in both root and rhizosphere compartments, with the host genotype explaining as much as 7.61% of the overall variance of the 16S rRNA, and 13.5% of ITS2 data (Fig. 3; P<0.001). The rhizosphere compartments of wild type and *ram1* were found to harbor similar bacterial communities, but were separate from those of *symrk* and *ccamk* (Fig. 3A). Further, the rhizosphere communities of each of these four plant genotypes were found to be significantly different from that of *nfr5* (Fig. 3A). A similar trend was observed for fungal communities, except that wild-type and *ram1* rhizosphere communities were clearly separated from each other (Fig. 3C). In the root compartment we found bacterial consortia that were distinctive for each of the five plant genotypes (Fig. 3B). This genotype effect was also found in the root-associated fungal communities, with the exception of *nfr5*, which was indistinguishable from wild-type (Fig. 3D). We then tested the contribution of AM and rhizobial symbionts to the observed patterns of diversity, in order to determine if AM fungi (Glomeromycota) and nitrogen-fixing *Mesorhizobium loti* (Phylobacteriaceae) are the sole drivers of these host genotype community shifts (Fig. 3). We performed an *in silico* experiment in which sequencing reads of these two symbiotic taxonomic groups were removed from the analyses. Although we observed a decrease in the percentage of variance explained by the host genotype (Fig. S3 compared to Fig. 3), overall patterns of β-diversity remained unaltered, suggesting that other community members besides root nodule and arbuscular mycorrhizal symbionts contribute to the plant genotype-specific community shifts. Collectively, our analyses of *L. japonicus* symbiotic mutants grown in natural soil show that lack of AM and/or RNS symbioses has a significant effect on plant growth and on the structures of bacterial and fungal communities associated with legume roots.

**Fig3.**
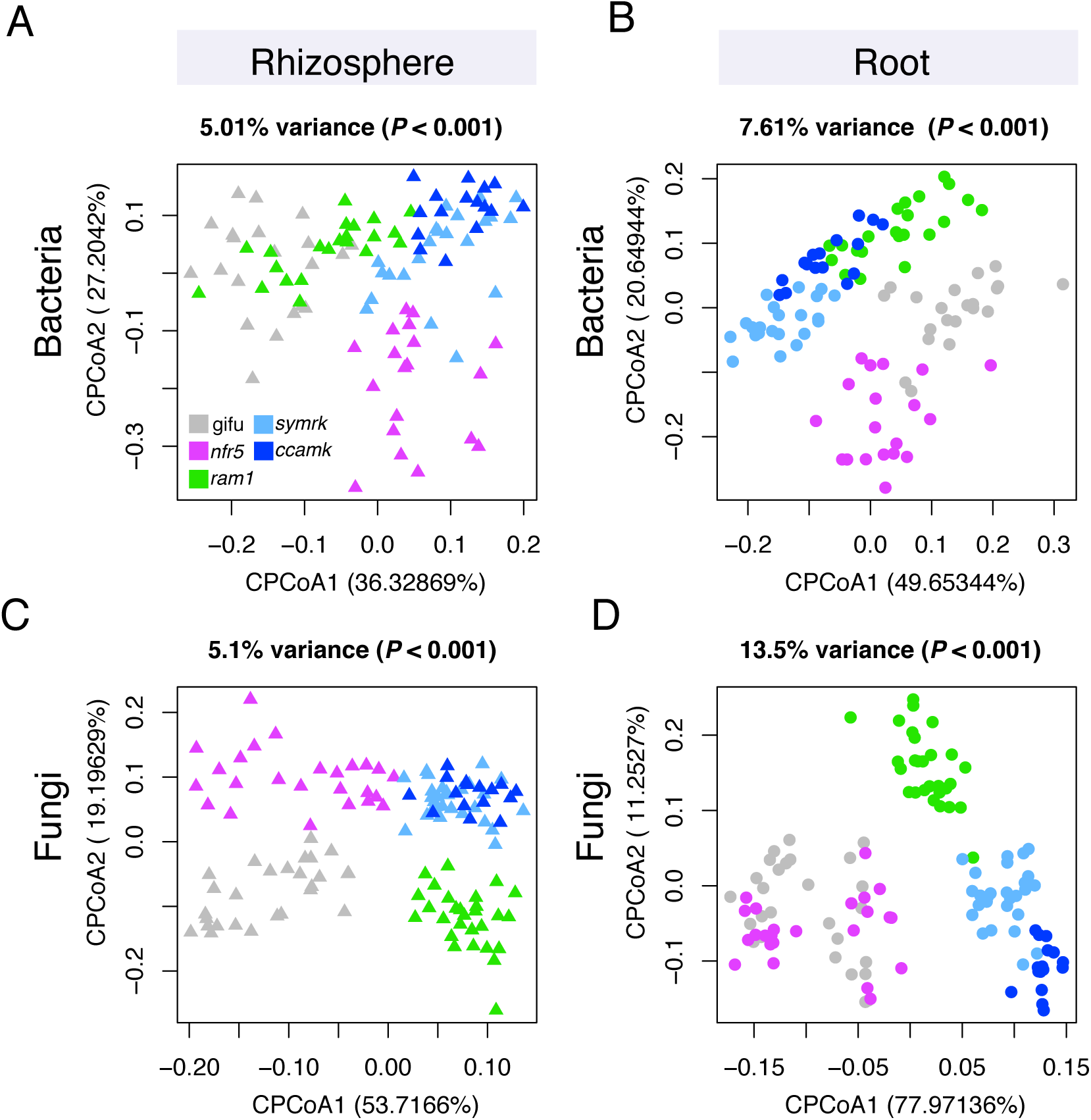
Constrained PCoA analysis showing genotype effect on microbial communities. A) Constrained PCoA plots for bacterial datasets showing rhizosphere samples (n = 100) and B) root samples (n = 100). C) Constrained PCoA plots for fungal datasets showing only rhizosphere samples (n = 124) and D) root samples (n = 122).

### Loss of symbiosis affects specific bacterial and fungal families of the root microbiota

Comparison of bacterial family abundances between wild type and mutants lacking RNS and/or AM symbiosis identified significant changes in Comamonadaceae, Phyllobacteriaceae, Methylophilaceae, Cytophagaceae and Sinobacteraceae in the rhizosphere compartment (Fig. 4A; top 10 most abundant families). The abundance of Comamonadaceae and Phyllobacteriaceae also differed significantly in the root compartment of RNS mutants compared to wild type. Streptomycetaceae and Sinobacteraceae were specifically affected by loss of *Nfr5*, whereas Anaeroplasmataceae and Burkholderiaceae were affected by the lack of AM symbiosis in *symrk* and *ccamk* plants (Fig. 4A). The relative abundances of the same two families were also significantly reduced in *ram1* roots, suggesting that active AM symbiosis influences root colonization by a subset of bacterial root microbiota taxa. Six out of the ten most abundant fungal families in the rhizosphere compartment of *Lotus* plants belonged to Ascomycota (Fig. 4B). By contrast, the root endosphere was dominated by numerous families of Glomeromycota, which were found to be almost fully depleted from the rhizosphere and root compartments of *ram1, symrk* and *ccamk* mutants, indicating that absence of AM symbiosis predominantly affects Glomeromycota and does not limit root colonization or rhizosphere association by other fungal families. However, depletion of Glomeromycota in the AM mutant roots was accompanied by an increase in the relative abundance of Ascomycota members belonging to Nectriaceae in both rhizosphere and root compartments and by an increased abundance of unclassified Helotiales, Leotiomycetes, and Sordariomycetes in the root compartment only (Fig. 4B).

**Fig4.**
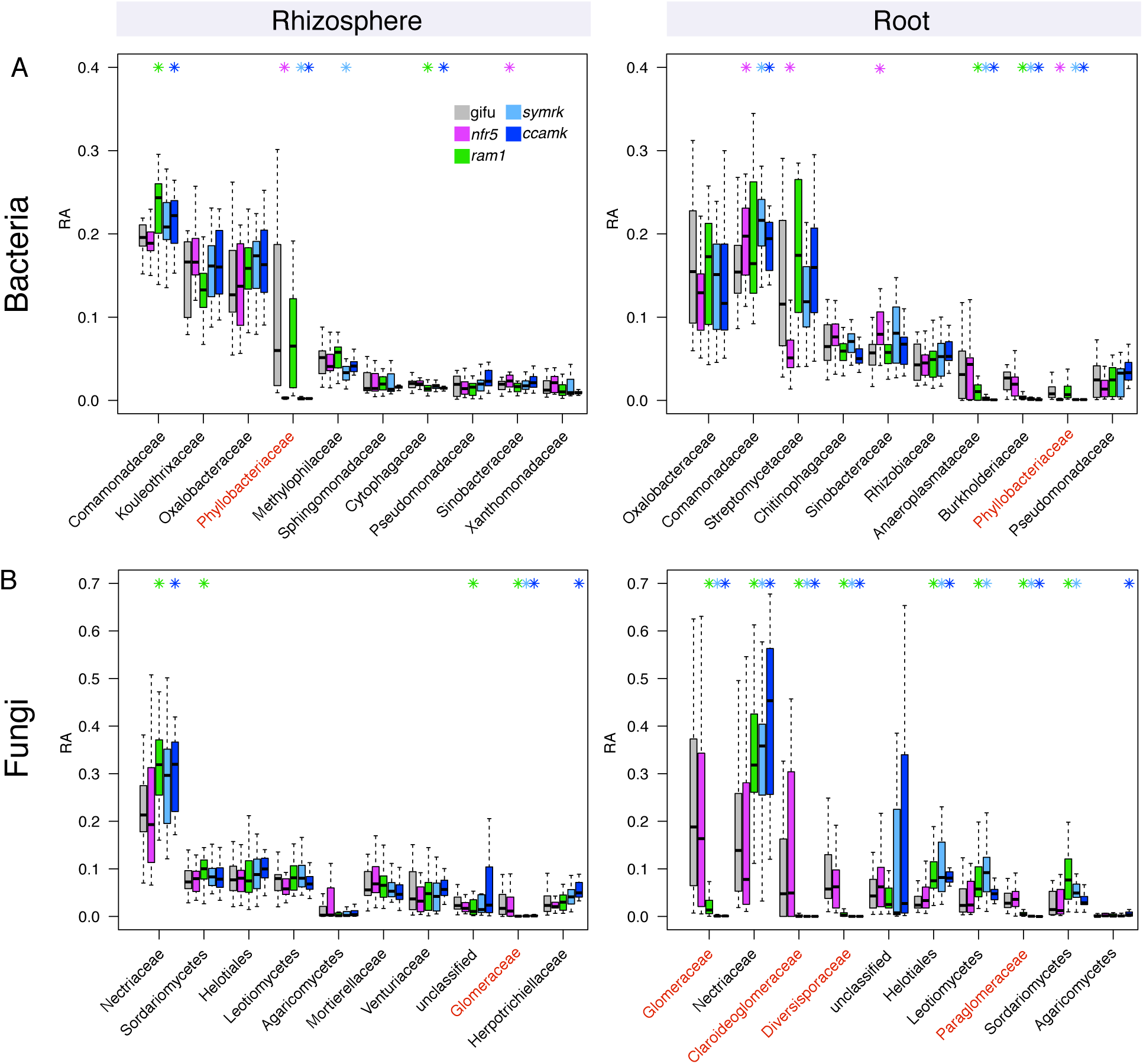
Relative abundance for main microbial taxa across plant compartments and genotypes. A) RA for bacterial families in rhizosphere (left panel) and root compartment (right panel). B) RA for fungal families in rhizosphere (left panel) and root compartment (right panel). Taxa are sorted in decreasing order according to their average RA in wt plants (only first 10 most abundant taxonomical groups are shown). RA in wt as well as in the respective mutants is displayed. Significant differences compared to wt are marked with an asterisk in the color of the mutant (P<0.05, Kruskal-Wallis test). Families that include known symbionts are marked in red (Phyllobacteriaceae for bacteria and Glomeromyctes for Fungi). For some fungal taxa the next higher rank is shown, when no family level information was available.

Closer inspection of the microbial community shifts at the OTU level identified 45 bacterial OTUs and 87 fungal OTUs enriched in the roots of symbiosis mutants compared to those of wild type (Fig. 5), and 60 bacterial OTUs and 30 differentially abundant fungal OTUs in the rhizosphere samples (Fig. S4). The absence of RNS in *nfr5* roots affected the relative abundance of multiple OTUs (n=27 in the root, n=23 in the rhizosphere) belonging to diverse taxa. Many of these OTUs (n=18 in the root, n=16 in the rhizosphere) showed a similar differential relative abundance in *symrk* and/or *ccamk* mutants when compared to wild type (Fig.5A), indicating that their contribution to the *Lotus* root communities outside of nodules is affected by active nitrogen fixing symbiosis. Impairment of both AM and RNS symbioses in *symrk* and/or *ccamk* mutants resulted in opposite changes in the relative root abundances of OTUs belonging to specific Burkholderiales families. Depletion of OTUs belonging to Burkholderiaceae (n=5) was accompanied by the enrichment of OTUs from other Burkholderiales families (Oxalobacteraceae [n=3], Comamonadadaceae [n=2], and Methylophilaceae [n=2]; Fig. 5A). Only three of the above-mentioned Burkholderiaceae OTUs were depleted in *ram1* roots, suggesting that their enrichment in *Lotus* roots is dependent on functional AM symbiosis.

**Fig5.**
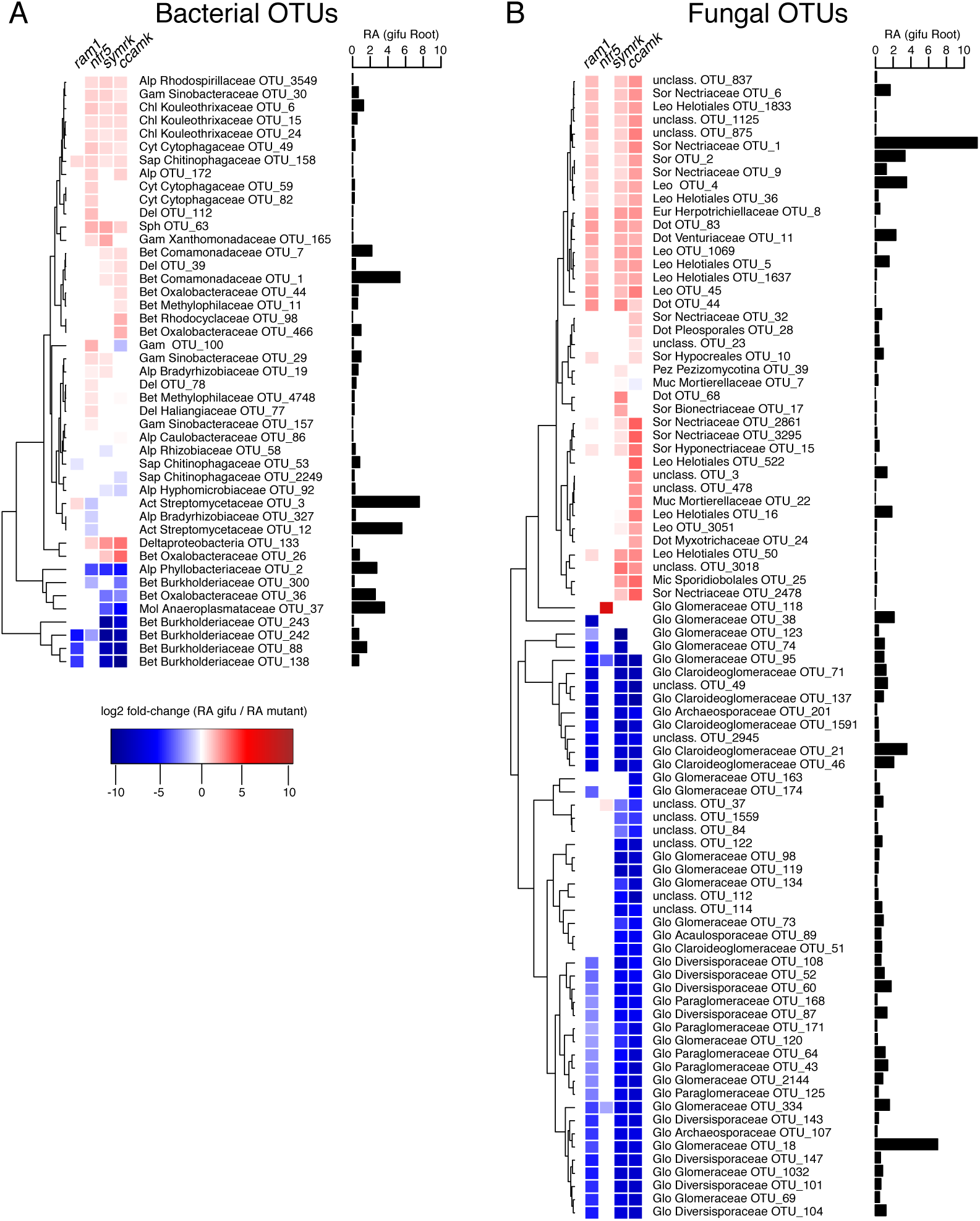
Differential abundance analysis for root associated OTUs. A) Bacterial OTUs that are differentially abundant in the roots of mutants compared to wt roots. B) fungal OTUs that are differentially abundant in the roots of mutants compared to wt roots. Only OTUs that have an average RA > 0.1% across all root samples, including mutants, are considered here. For each OTU the fold change in RA from wt to mutant is indicated (P <0.05, Kruskal-Wallis test). Next to each OTU the RA in wt roots is indicated. Phylum and family asscociation (if available) is given for each OTU (Bacterial phyla: Del=Deltaproteobactria, Gem=Gemm-1, Chl=Chloroflexi, Bet=Betaproteobacteria, Alp=Alphaproteobacteria, Gam=Gammaproteobacteria, Cyt=Cytophagia, Sap=Saprospirae, Ped=Pedosphaerae, Sph= Sphingobacteria, Mol= Mollicutes; Fungal phyla: Sor=Sordariomyctes, Dot=Dothideomycetes, Mic= Microbotryomycetes, Ust=Ustilaginomycetes, Eur=eurotiomycetes, Leo=Leotiomycetes, Aga=Agaricomycetes, Glo=Glomeromyctes, Pez=Pezizomycota, Muc=Mucoromycotina).

Analysis of the ITS2 amplicon sequences from root samples identified a large number of Glomeromycota OTUs (n=39), demonstrating the capacity of *Lotus* Gifu roots grown in natural soil to accommodate a phylogenetically diverse community of AM fungi (Fig. 5B). The majority of these fungal OTUs (n=31) were depleted in *symrk, ccamk* and *ram1* mutant roots, indicating that their enrichment is dependent on a functional AM symbiosis pathway. Their intraradical colonization appears to be independent of *RAM1*, as 12 OTUs assigned to Glomeromycota or unclassified, nine of which define a Glomeromycota sublineage, were depleted in *symrk* and *ccamk* but not in *ram1* roots. A reduced abundance of Glomeromycota OTUs in the endosphere compartment was accompanied by an increased abundance of Ascomycota members, especially of members belonging to the Nectriaceae (8 OTUs) and Helotiales (7 OTUs) families, suggestive of a mutually exclusive occupancy of the intraradical niche. In sum, our results reveal that in natural soil CSSP symbiotic genes are essential for root colonization by a wide range of Glomeromycota fungi and that these genes significantly affect the abundances of multiple bacterial taxa, predominantly belonging to the Burkholderiales and Rhizobiales orders.

In order to assess the impact of mutations of *Lotus* symbiotic genes on microbial interactions we constructed co-occurrence microbial networks for each genotype independently using SparCC [37] (Fig. S5). We observed an increase in the number of edges of the networks inferred from *symrk* and *ccamk* (748 and 805 edges, respectively) compared to Gifu WT, *nfr5*, and *ram1* networks (471, 569 and 500 edges, respectively; Fig S5 A), despite a comparable number of nodes in all genotypes. This unexpected observation suggests a greater connectivity between bacterial root commensals when both fungal and bacterial symbioses are disrupted in *symrk* and *ccamk* roots. In the corresponding five fungal networks the number of OTUs is moderately reduced in *ram1* and approximately halved in *symrk* and *ccamk* networks (86 in Gifu WT, 78 in *nfr5*, 63 in *ram1*, 39 in *symrk*, 41 in *ccamk*; Fig S5 A), which can be explained by the partial or complete depletion of Glomeromycota taxa in the latter three host genotypes. This decrease in the number of fungal OTUs is accompanied by a decrease in the number of edges in the fungal networks (329 Gifu, 363 *nfr5*, 231 *ram1*, 101 *symrk* and 117 edges in *ccamk*, respectively; Fig S5 A). To directly compare the number of edges between plant genotypes for bacterial and fungal networks we first normalized the number of bacterial and fungal OTUs (Fig S5 B). Compared to Gifu WT and *nfr5* networks the degree centrality for bacterial OTUs is slightly increased in *ram1* (significant only for positive correlations) and clearly increased in *symrk* and *ccamk* (significant for both positive and negative correlations), supporting the aforementioned change in network structure of the bacterial root microbiota when both fungal and bacterial symbiosis are disrupted in *Lotus* roots. By contrast, the degree centrality of fungal OTUs remains mostly stable across fungal networks identified in the five plant genotypes. Together our analyses suggest that the combined activities of fungal and bacterial symbioses negatively influence the connectivity within the *Lotus* bacterial root microbiota.

## Discussion

Here we investigated the role of host AM and/or RNS genes in establishing structured bacterial and fungal communities in the rhizosphere and endosphere compartments of *L. japonicus* grown in natural soil. Impairment of RNS in *nfr5* or AMS in *ram1* plants had a significant impact on root microbiota structure, which was mainly, but not exclusively, confined to the composition of corresponding bacterial or fungal communities, respectively (Fig. 3, 4, and 5).

The shift between the root-associated microbial communities of wild type and *nfr5-2* mutant is in line with both the qualitative and quantitative findings of a previous report on the *Lotus* bacterial root microbiota (Fig. 3A and B in this study; [22]). Here, however, we observed an enhanced rhizosphere effect in both wild type and *nfr5* plants, leading also to a less prominent community shift in this compartment (Fig. S6), which was not previously observed. These differences in rhizosphere bacterial composition are likely caused by a soil batch effect and, to a lesser extent, possibly also the use of different sequencing platforms (Illumina in this study *versus* 454 pyrosequencing in [22]). The nearly unaltered fungal community composition in *nfr5* mutant plants compared to wild type (only 3 out of 39 Glomeromycota OTUs differentially abundant) suggests that NFR5 is dispensable for fungal colonization of *L. japonicus* roots. This is consistent with recent findings from analyses of diverse AM symbiotic mutants of *Lotus* where the structure of the root-associated fungal communities of AM- and CSSP-deficient mutants was indistinguishable [25]. Despite unaltered fungal communities in *nfr5* mutants, we found a marked shoot biomass reduction of this genotype grown in natural soil (∼4-fold; Fig. 2), revealing that intraradical colonization by soil-derived fungal endophytes is robust against major differences in plant growth.

A recent microbial multi-kingdom interaction study in *A. thaliana* showed that bacterial commensals of the root microbiota are crucial for the growth of a taxonomically wide range of fungal root endophytes. These antagonistic interactions between bacterial and fungal root endophytes are essential for plant survival in natural soil [32]. We have shown here that an almost complete depletion of diverse Glomeromycota taxa from roots of each of the three AM mutants was accompanied by an enrichment of fungal OTUs belonging to the families Nectriaceae and Helotiales (Fig. 4). We speculate that the increased relative abundance of these fungal taxa is caused by intraradical niche replacement as a compensatory effect following the exclusion of Glomeromycota symbionts from the root compartment. Previous mono-association experiments have shown that isolates belonging to Nectriaceae and Helotiales can have either mutualistic or pathogenic phenotypes [38,39,40]. Given that all plant genotypes were free of disease symptoms when grown in natural soil (Fig. 2), we speculate that the complex shifts in the composition of the bacterial root microbiota in *nfr5, symrk*, and *ccamk* mutants did not affect the capacity of bacterial endophytes to prevent pathogenic fungal overgrowth. Of note, Helotiales root endophytes were also enriched in roots of healthy *Arabis alpina*, a non-mycorrhizal plant species and relative of *A. thaliana*, and contribute to phosphorus nutrition of the host when grown in extremely phosphorus-impoverished soil [41]. The enrichment of Helotiales in *Lotus* AM mutants is therefore consistent with potential niche replacement by other fungal lineages to ensure plant nutrition in nutrient-impoverished soils. Although the proposed compensatory effect in AM mutants will need further experimental testing in phosphorus-depleted soils, our hypothesis is consistent with the only mild impairment in plant growth in *ram1* mutants (Fig. 2).

We observed that members of the bacterial families of Burkholderiaceae and Anaeroplasmataceae are significantly depleted in the roots of each of the three AM mutants compared to wild type. Members of the Glomeromycota have been found to contain intracellular endosymbiotic bacteria [42], some belonging to the order Burkholderiales [41]. Interestingly, the most positively correlated bacterial OTUs with Glomeromycota fungi in our network analyses included one Anaeroplasmataceae and two Burkholderiaceae OTUs (Fig. S7), further indicating a direct interaction between these taxonomic groups. These findings suggest that these bacteria are either endosymbionts of Glomeromycota fungi that are excluded from the roots of the AM defective genotypes or that their intraradical colonization is indirectly mediated by AM fungus infection. Except small changes in the bacterial root microbiota in *ram1* plants, which are mainly limited to the aforementioned Burkholderiaceae and Anaeroplasmataceae OTUs, the structure of the root-associated bacterial community is remarkably robust against major changes in the composition of root-associated fungal assemblages (Fig. 5). Nevertheless, we observed a clear increase in connectivity between bacterial OTUs and degree centrality parameters in the bacterial networks constructed from *symrk* and *ccamk* mutants compared to those of Gifu, *nfr5* and *ram1*. This unexpected change in bacterial network structure could be a consequence of a vacant niche created by depletion of dominant Glomeromycota taxa from the interior of *symrk* and *ccamk* roots. But niche filling by bacterial commensals is unlikely to explain the observed alteration in bacterial network connectivity because Glomeromycota root colonization is greatly diminished in *ram1* plants without major changes in the corresponding bacterial network structure (Fig. 4 and Fig. S5). Increased bacterial network connectivity in *symrk* and *ccamk* roots is more likely a consequence of inactivation of the CSSP, which remains intact in all other tested genotypes. However, we cannot fully exclude that the altered nutritional status in *symrk* and *ccamk* plants resulting from the combined loss of metabolic activities of, and induced by both symbionts also plays a role in the altered network structure.

Paleontological and phylogenomic studies established the ancestral origin of genetic signatures enabling AM symbiosis in land plants [1,44]. In monocots and dicots, the extended AM fungal network is primarily recognized as a provider of nutrients, particularly phosphorus [45,46], but the positive impact of AM symbiosis on the host transcends nutrient acquisition [47]. Additionally, phylogenomic studies of the symbiotic phosphate transporter PT4 suggest that this trait evolved late and therefore that phosphorus acquisition might not have been the (only) driving force for the emergence of AM symbiosis [44]. *SymRK* and *Ram1* were identified in the genomes of liverworts, but evolution of *CCaMK* predated the emergence of all land plants, as shown by its presence and conserved biochemical function in advanced charophytes [44]. Together, these findings raise questions regarding the forces driving the evolution of signaling genes enabling intracellular symbioses in land plants. Our study shows that in *L. japonicus*, simultaneous impairment of AM and RN symbioses in *symrk* and *ccamk* plants had a dramatic effect on the composition of both bacterial and fungal communities of the legume root microbiota (Fig. 5). Importantly, mutation of *CCaMK* and *SymRK* led to an almost complete depletion of a large number of fungal OTUs, mostly belonging to Glomeromycota, indicating that in *Lotus*, these genes predominantly control the colonization of roots by this particular fungal lineage. The finding that *ram1-2* mutants show retained accommodation for a subset of fungal root endophytes (n=13; Fig. 5B, and Fig 4B) whose colonization is dependent on an intact common symbiosis pathway is not surprising based on the capacity of these mutants to enable fungal colonization but not to sustain a full symbiotic association [35], and indicates that *RAM1* is dispensable for the intraradical colonization of these Glomeromycota fungi. Alternatively, these fungal root endophytes may engage in commensal rather than mutualistic relationships with *L. japonicus* independently of the AM symbiosis pathway, as is the case for multiple species of commensal non-symbiotic rhizobia [22,48]. Given that *ram1* mutants specifically block AM arbuscule differentiation but not root colonization [35], it is conceivable that the Glomeromycota taxa colonizing this plant genotype cannot form arbuscules during root colonization.

Legumes have evolved the capacity to recognize and accommodate both types of intracellular symbionts, and the large effect of CSSP genes on associated microbiota seen in the present work could reflect a legume-specific trait. However, in rice, which does not engage in symbiotic relationships with nodulating rhizobia, mutants lacking *CCaMK* were also found to display significant changes in root-associated bacterial communities that could be mainly explained by depletion of Rhizobiales and Sphingomonadales lineages [49]. Thus, our findings based on comparative microbiota analysis of *Lotus ccamk* and *ram1* mutants suggest a broader role for common symbiosis signaling genes in microbiota assembly. Future studies on orthologous genes in basal land plants will contribute to a better understanding of the role of symbiotic signaling in the evolution of plant-microbiota associations.

## Materials and Methods

### Preparation and storage of soil

The two soil batches used in this study were collected from Max Planck Institute for Plant Breeding Research agricultural field located in Cologne, Germany (50.958N, 6.865E) in the following seasons: CAS11-spring/autumn 2016, CAS12-spring 2017 (CAS: Cologne Agriculture Soil). The field had not been cultivated in previous years, no fertilizer or pesticide administration took place at the harvesting site. Following harvest, soil was sieved, homogenized and stored at 4 °C for further use.

### Soil and plant material

All studied *L. japonicus* symbiosis-deficient mutants, *nfr5-2* [33], *ram1-2* [35], *symrk-3* [7] and *ccamk-13* [34], originated from the Gifu B-129 genotype.

### Plant growth and harvesting procedure

The germination procedure of *L. japonicus* seeds included sandpaper scarification, surface sterilisation in 1% hypochlorite bleach (20 min, 60 rpm), followed by three washes with sterile water and incubation on wet filter paper in Petri dishes for one week (temperature: 20 °C, day/night cycle 16/8h, relative humidity: 60%). For each genotype and soil batch, six to eight biological replicas were prepared by potting four plants in 7×7×9 cm pot filled with corresponding batch of soil (CAS11 soil six replicates, CAS12 soil eight replicated). For each batch of soil, two independent experiments have been carried out. Plants were incubated for ten weeks in the greenhouse (day/night cycle 16/8h, light intensity 6000 LUX, temperature: 20 °C, relative humidity: 60%), and were watered with tap water twice per week.

The block of soil containing plant roots was removed from the pot and adhering soil was discarded manually. Three sample pools were collected: complete root systems (harvested 1 cm below the hypocotyl), upper fragments of the root systems (4 cm-long, starting 1 cm below the hypocotyl) and lower root system fragments (harvested from 9 cm below; the latter two were collected from plants grown in the same pot (Fig. 1a). All pools were washed twice with sterile water containing 0.02% Triton X-1000 detergent and twice with pure sterile water by vigorous shaking for 1 min. The rhizosphere compartment was derived by collection of pellet following centrifugation of the first wash solution for 10 min at 1500 g. The nodules and visible primordia were separated from washed root pools of nodulating genotypes (WT and ram1-2) with a scalpel and discarded. In order to obtain the root compartment the root sample pools were sonicated to deplete the microbiota fraction attached to the root surface. It included 10 cycles of 30-second ultrasound treatment (Bioruptor NextGen UCD-300, Diagenode) for complete root systems and upper root fragments, while for the lower root fragments the number of cycles was reduced to three. All samples were stored at −80 °C for further processing. For AM colonisation inspection the whole root system of washed soil-grown plants was stained with 5% ink in 5% acetic acid solution and inspected for intraradical infection.

### Generation of 16S rRNA and ITS2 fragment amplicon libraries for Illumina MiSeq sequencing

Root pool samples were homogenized by grinding in a mortar filled with liquid nitrogen and treatment with Precellys24 Tissue lyser (Bertin Technologies) for two cycles at 5600 rpm for 30 sec. DNA was extracted with the FastDNA Spin Kit for Soil, according to the manufacturer’s protocol (MP Bioproducts). DNA concentrations were measured fluorometrically (Quant-iT™ PicoGreen dsDNA assay kit, Life Technologies, Darmstadt, Germany), and adjusted to 3.5 ng/μl. Barcoded primers targeting the variable V5-V7 region of the bacterial 16S rRNA gene (799F and 1193R, [29]) or targeting the ITS2 region of the eukaryotic ribosome (fITS7 and ITS4, [30,31]) were used for amplification. The amplification products were purified, pooled and subjected to sequencing with Illumina MiSeq equipment.

### Processing of 16S rRNA and ITS2 reads

Libraries from the three root fractions (including root tips endosphere, upper root endosphere and whole root endosphere) were analysed independently. As well as the main experiments (only whole roots). Due to a very low read count for 16S data in the first experiment in CAS11 soil, this data was not included in the final analysis. This resulted in an overall lower sample number for bacteria than for fungi (222 vs. 274 samples). All sets of amplicon reads were processed as recently described [32], using a combination of QIIME [50] and USEARCH tools [51]. For both datasets paired end reads were used. For ITS2 data, forward reads were kept, in case that no paired version was available. Main steps include quality filtering of reads, de-replicating, chimera detection and OTU clustering at a 97% threshold. 16S reads were filtered against the greengenes data base [52], whereas for ITS2 the reads where checked with ITSx [53] and compared against a dedicated ITS database to remove ITS sequences from non-fungal species. Taxonomic classification was done with uclust (assign_taxonomy from QIIME) for 16S OTUs and rdp classifier [54] for ITS2 OTUs. For the sake of consistency with NCBI taxonomic classification, the assignment of the ITS2 sequences was manually corrected so that that all OTUs assigned as Ilyonectria were assigned as belonging to the Sordariomycetes, Hypocreales and Nectriaceae. For 16S data, OTUs assigned as mitochondrial or chloroplast, were removed prior to analysis.

### Statistical analysis

For calculating Shannon diversity indices, OTU tables were rarefied to 1000 reads (single_rarefaction.py from QIIME, samples with less than 1000 reads were omitted). Significant differences were determined using ANOVA (aov function in R) and post-hoc Tukey test (TukeyHSD in R, p<0.05). For calculating Bray Curtis distances between samples, OTU tables were normalized using cumulative sum scaling (CSS, [55]). Bray-Curtis distances were used as input for principal coordinate analysis (PCoA, cmdscale function in R) plots and as input for constrained analysis of principal coordinate (CPCoA, capscale function, vegan package in R). For the latter, the analysis was constrained by genotypes (each mutant and WT separately) and corrected for the effect of the two soil types (CAS 11, CAS 12) and the four individual experiments (using the “Condition” function). This analysis has been repeated with OTU tables from which OTUs that represent known plant symbionts (Phyllobacteriaceae for 16S and Glomeromycota for ITS2) were removed before normalization, distance calculation and CPCoA. A previously described approach was used to draw ternary plots and for respective enrichment analysis [22]. Fold change of OTUs between wild type and mutant plants was calculated as followed. Samples showing a read count <5000 were removed. OTUs with mean relative abundance (RA) >0.1% across all root or rhizosphere samples, respectively, were kept for analysis. Fold change in RA from WT to mutants was calculated over all WT samples for *nfr5, ram1* and *symrk* whereas the change to *ccamk* was only calculated with WT samples from experiments where *ccamk* mutants were present. To avoid zeros in calculation the RA of OTUs missing from samples was set to 0.001%. The significance of differences in abundance was tested using the Kruskal-Wallis test (p<0.05). Networks for each genotype and kingdom were calculated independently using SparCC [37]. OTU tables were filtered before analysis to only include samples from one soil type (CAS12) to avoid biases. In addition only OTUs that are present in more than 10 samples and have mean RA >0.1% were kept for network analysis. Raw count tables were given to SparCC as an input, the resulting correlations were filtered by significance (p < 0.05). Networks were drawn using Cytoscape [56]. To calculate the degree centrality, the number of positive and negative connections, for each OTU was divided by the number of OTUs present in the respective network. Correlations between bacterial and fungal OTUs were calculated as followed. OTUs that appear in less than 10 gifu root samples and have mean RA < 0.1% were not considered for this analysis. Spearman rank correlations were calculated between RA values of bacterial and fungal OTUs across all gifu root samples (cor.test function in R, p< 0.001). For showing the cumulative correlation of bacterial OTUs to fungal OTUs, the respective correlations for one bacterial OTU were summed up, so the number of correlations and the strength could be accessed in one analysis. This was repeated, but just for fungal OTUs annotated as Glomeromycota.

## Acknowledgements

This work was supported by funds to S. R. from The Danish National Research Foundation (Grant no. DNRF79), by funds to P.S.-L. from the Max Planck Society, a European Research Council advanced grant (ROOTMICROBIOTA), the ‘Cluster of Excellence on Plant Sciences’ program funded by the Deutsche Forschungsgemeinschaft (DFG), and SPP 2125 DECRyPT from the DFG.

## Data availability

All sequencing reads will be uploaded to the European Nucleotide Archive (ENA). Code and relevant data files (e.g. OTU tables) will be made public via GitHub.

## Supplementary Figures

**supp Fig1.**
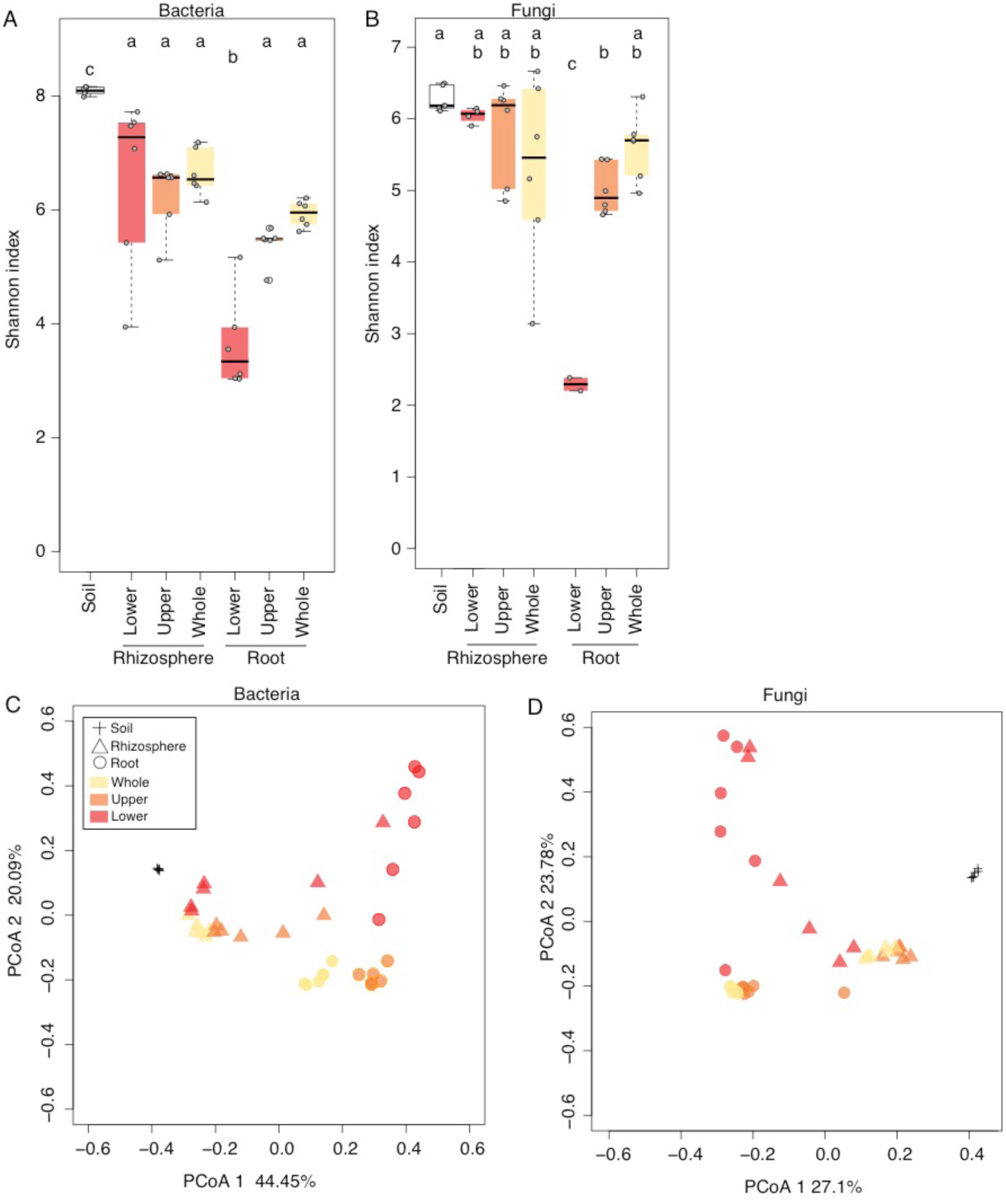
Alpha and beta diversity across root fractions. A) Shannon diversity index for 16S amplicon data, soil (n=6), lower (n=6), upper (n=6), whole root fractions (n=6), and respective rhizosphere samples (n=6 each) B) Shannon diversity index for ITS2 amplicon data, soil (n=6), lower (n=2), upper (n=6), whole root fractions (n=6), and respective rhizosphere samples (n=6 each, except lower, n=4) (ANOVA with Tukey’s post hoc test, P<0.05). C) Principal coordinate analysis of Bray-Curtis distances for bacterial data. D) Principal coordinate analysis of Bray-Curtis distances for fungal data.

**supp Fig2.**
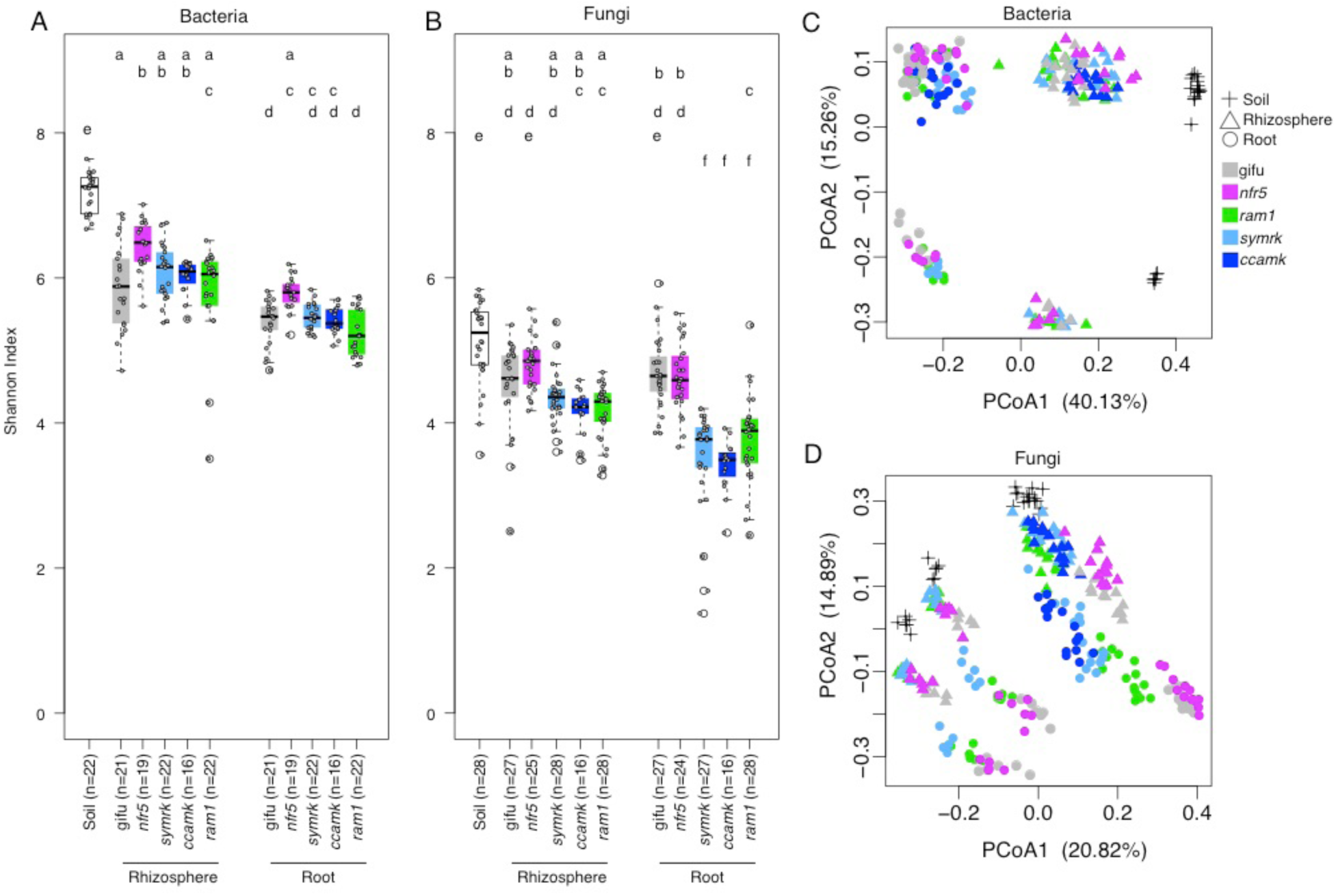
Alpha and beta diversity across plant compartments and genotypes. A) Shannon diversity indices for the bacterial (16S amplicon) dataset. B) Shannon diversity indices for the fungal (ITS2 amplicon) dataset (ANOVA with Tukey’s post hoc test, P<0.05) C) Principal coordinate analysis of Bray-Curtis distances for the bacterial dataset (n=222). D) Principal coordinate analysis of Bray-Curtis distances for the fungal dataset (n=274).

**supp Fig3.**
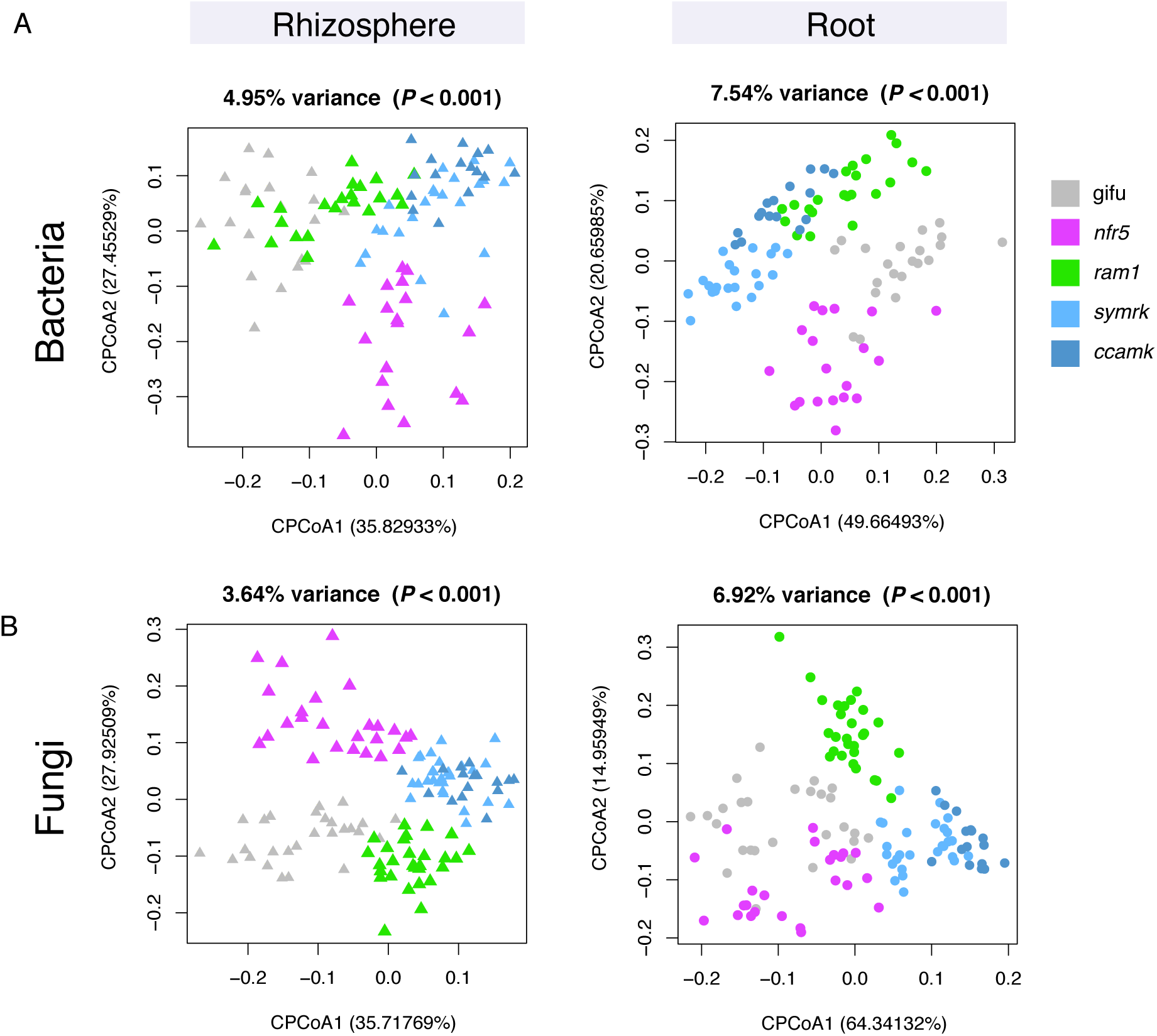
CPCoA with in-silico depletion of known symbionts. Results are separated by compartments. Datasets were constrained by genotype, and filtered for effects of experiments and soil type. A) Bacterial dataset from which OTUs belonging to the Phyllobacteriaceae were removed before analysis (root n=100, rhizosphere n=100). B) Fungal dataset from which OTUs belonging to the Glomeromycota were removed before analysis (root n=122, rhizosphere n=124).

**supp Fig4.**
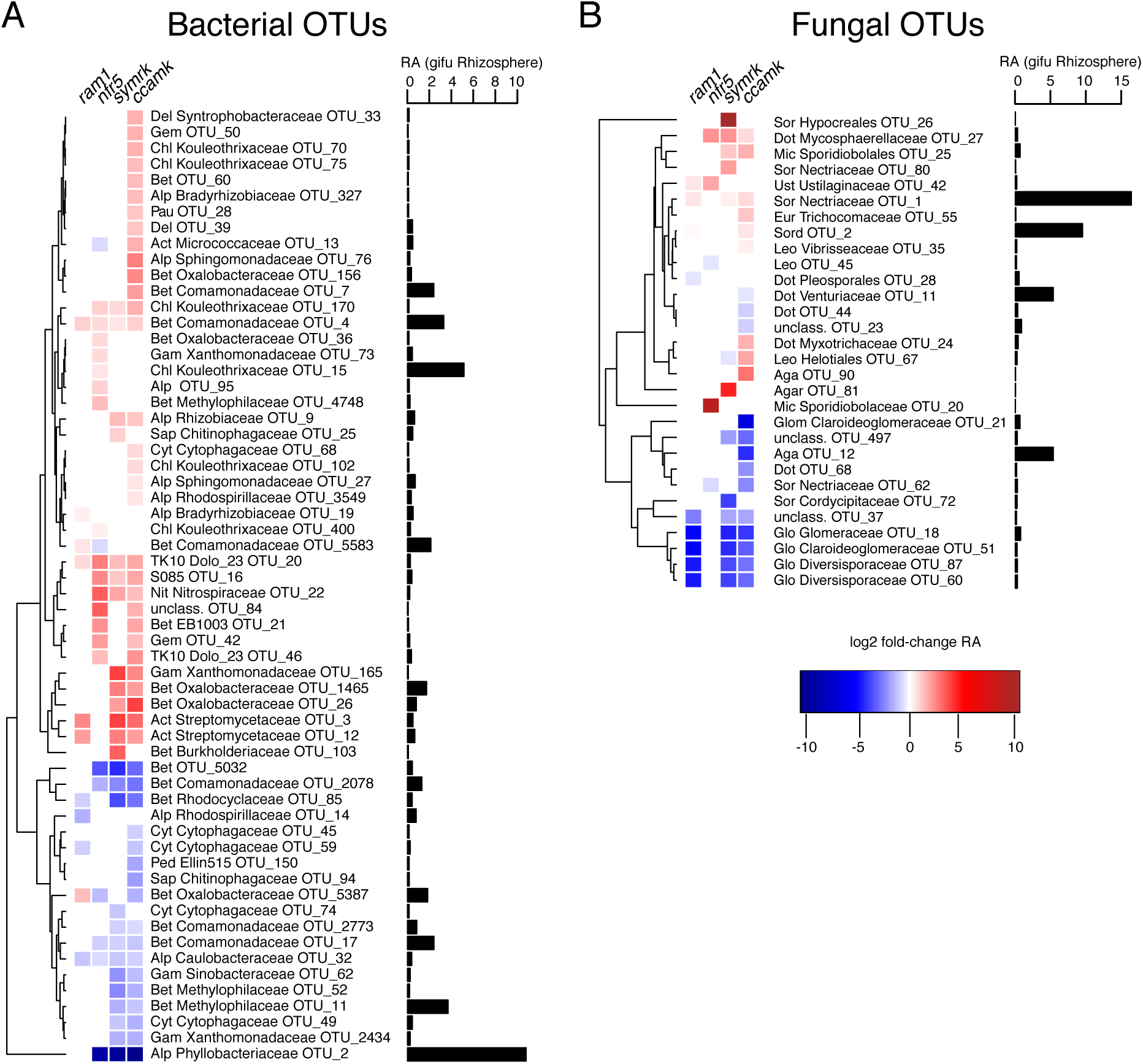
Differential abundance analysis for rhizosphere-associated OTUs showing enrichment and depletion in mutants. A) Bacterial OTUs that are differentially abundant in the rhizosphere of mutants compared to wt. B) Fungal OTUs that are differentially abundant in the rhizosphere of mutants compared to wt. Only OTUs that have an average RA > 0.1% across all rhizosphere samples, including mutants, are considered here. For each OTU, the fold change in RA from wt to mutant is indicated (P <0.05, Kruskal-Wallis test). Next to each OTU the RA in wt rhizosphere is indicated. Phylum and family asscociation is given for each OTU (Bacterial phyla: Del=Deltaproteobacteria, Gem=Gemm-1, Chl=Chloroflexi, Bet=Betaproteobacteria, Alp=Alphaproteobacteria, Gam=Gammaproteobacteria, Cyt=Cytophagia, Sap=Saprospirae, Ped=Pedosphaerae, Sph= Sphingobacteria, Mol= Mollicutes; Fungal phyla: Sor=Sordariomyctes, Dot=Dothideomycetes, Mic= Microbotryomycetes, Ust=Ustilaginomycetes, Eur=Eurotiomycetes, Leo=Leotiomycetes, Aga=Agaricomycetes, Glo=Glomeromyctes, Pez=Pezizomycota, Muc=Mucoromycotina).

**supp Fig5.**
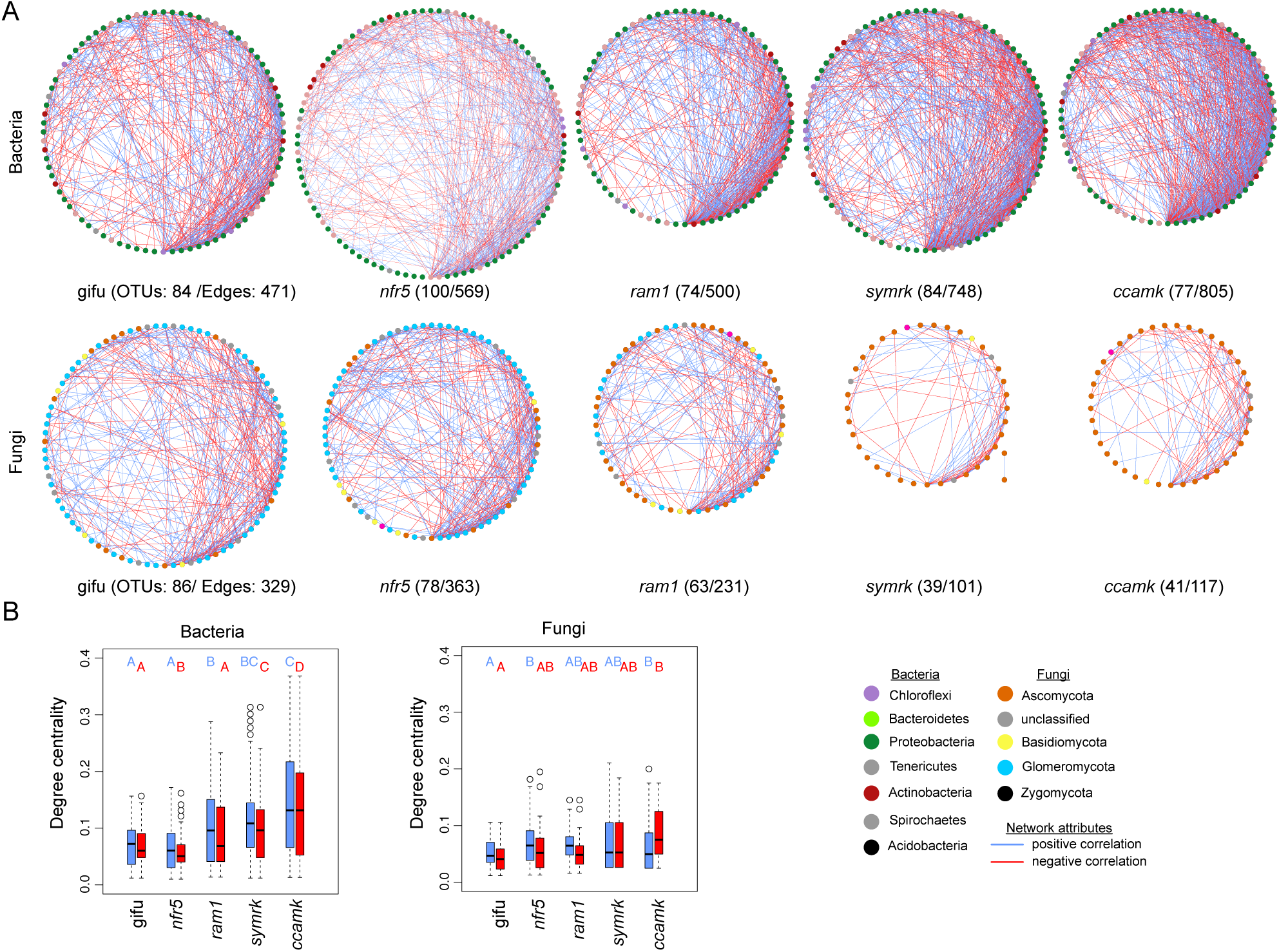
Network analysis of root-associated bacterial and fungal OTUs. A) SparCC based networks for wildtype and mutant roots (upper panel: bacterial OTUs, lower panel: fungal OTUs). Nodes are colored by phylum and are ordered by increasing degree (clockwise, degree sorted circle layout from Cytoscape). Positive connections are colored in blue, negative connections are colored in red. All connections are significant (SparCC, pseudo p-values p<0.05). B) Boxplots showing degree centrality for the different networks. (left panel: bacteria, right panel: fungi). Degree centrality was calculated separately for positive and negative edges, as well as the statistical test were done separate (pairwise Wilcox test, FDR <0.05).

**supp Fig6.**
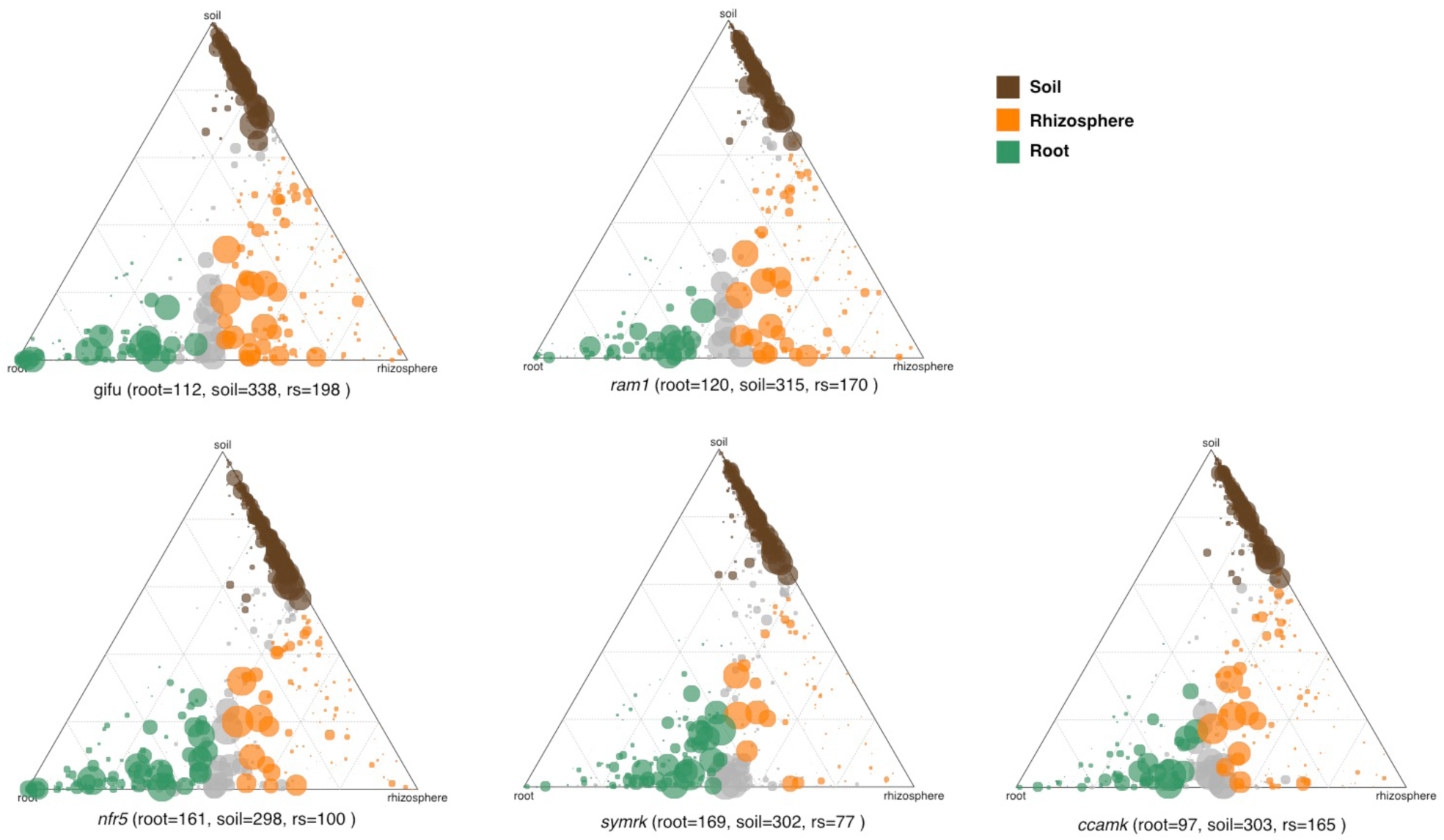
Ternary plots showing compartment-enriched bacterial OTUs. Separately for wt and mutant plants. Below each plot the number of enriched OTUs for each compartment is shown.

**supp Fig7.**
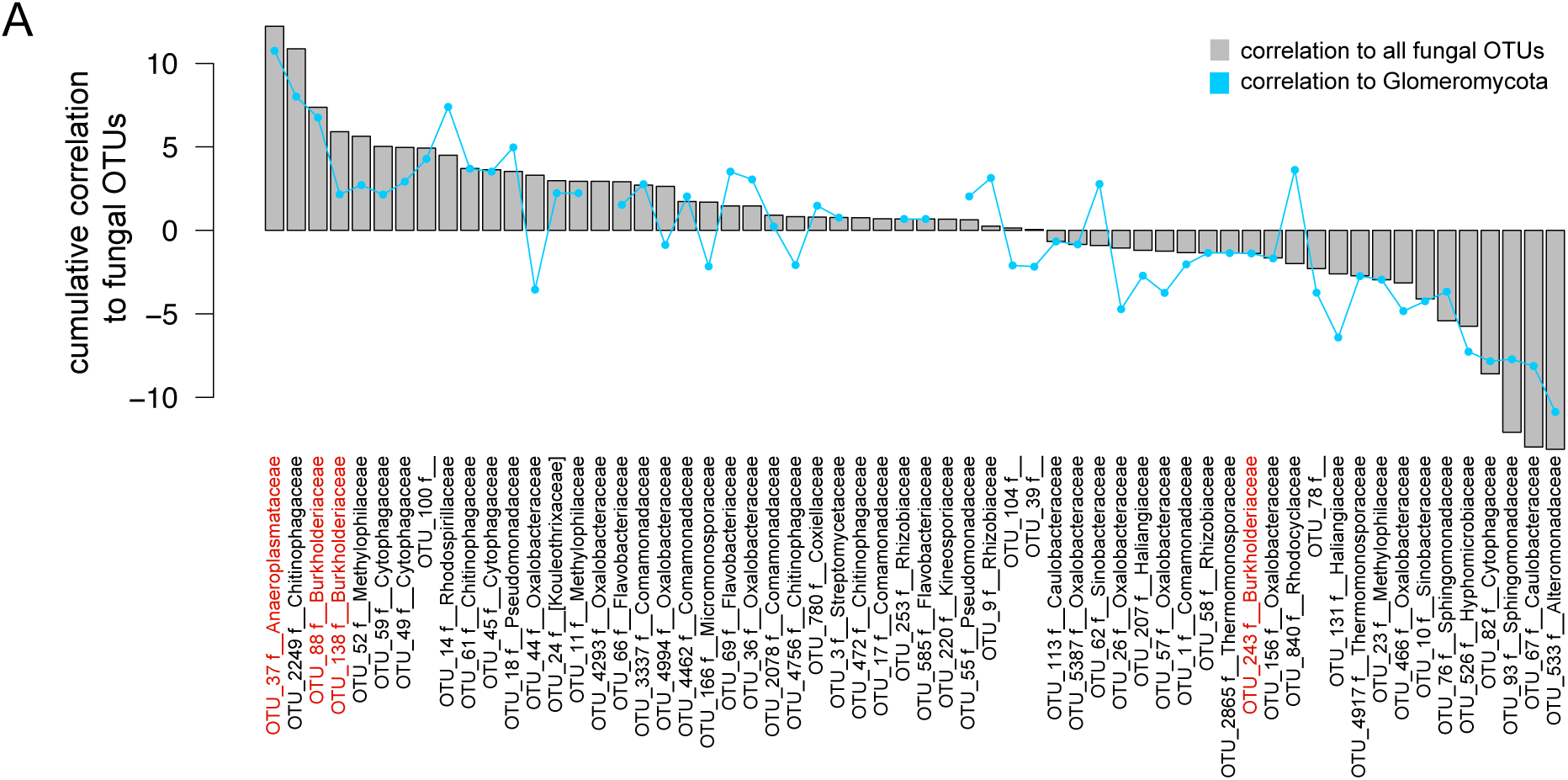
Interkingdom correlation of bacterial OTUs in wild-type roots. A) Bacterial OTUs that show a significant correlation to fungal OTUs across wild-type root samples (OTU RA >0.01%, spearman rank correlation p <0.001) are sorted according to their cumulated correlation to all fungal OTUs (grey bars). In addition the cumulative correlation only to Glomeromycotal OTUs is indicated by blue dots within each bar. Taxonomy at the family levels is given for each OTU, if available. OTUs belonging to the Burkholdericeae and Anaeroplasmataceae are highlighted in red.

**supp table1.**
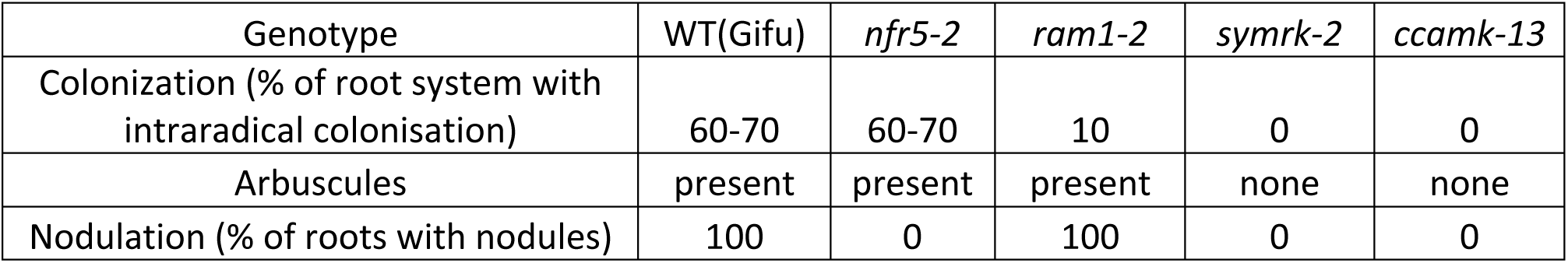
Symbiotic phenotype of *Lotus japonicus* wild-type and mutants grown in Cologne soil (*n=5*)

| Genotype | WT(Gifu) | *nfr5-2* | *ram1-2* | *symrk-2* | *ccamk-13* |
| --- | --- | --- | --- | --- | --- |
| Colonization (% of root system with intraradical colonisation) | 60-70 | 60-70 | 10 | 0 | 0 |
| Arbuscules | present | present | present | none | none |
| Nodulation (% of roots with nodules) | 100 | 0 | 100 | 0 | 0 |

